# Hyperspectral Segmentation of Plants in Fabricated Ecosystems

**DOI:** 10.1101/2024.12.20.629718

**Authors:** Petrus H. Zwart, Peter Andeer, Trent Northen

## Abstract

Hyperspectral imaging provides a powerful tool for analyzing above-ground plant characteristics in fabricated ecosystems, offering rich spectral information across diverse wavelengths. This study presents an efficient workflow for hyperspectral data segmentation and subsequent data analytics, minimizing the need for user annotation through the use of ensembles of sparse mixed-scale convolution neural networks. The segmentation process leverages the diversity of ensembles to achieve high accuracy with minimal labeled data, reducing labor-intensive annotation efforts. To further enhance robustness, we incorporate image alignment techniques to address spatial variability in the dataset. Down-stream analysis focuses on using the segmented data for processing spectral data, enabling monitoring of plant health. This approach not only provides a scalable solution for spectral segmentation but also facilitates actionable insights into plant conditions in complex, controlled environments. Our results demonstrate the utility of combining advanced machine learning techniques with hyperspectral analytics for high-throughput plant monitoring.

## 1 Introduction

EcoFABs, or fabricated ecosystems, are innovative laboratory devices designed to advance plant microbiome research by providing standardized and reproducible platforms for controlled experiments. These compact growth chambers offer precise control over environmental variables such as nutrient availability, light conditions, and microbial communities, enabling researchers to study the dynamics of plant growth and root-microbe interactions in a semi-high-throughput manner. By allowing spatially defined simultaneous imaging of root and shoot systems, EcoFABs facilitate comprehensive analysis of plant growth above and below ground. Their design also supports flexible sampling and the precise addition of microbes or materials, making them a versatile tool for fundamental plant science, sustainable agriculture, biosecurity, and applications such as nitrogen fixation, carbon sequestration, and even exobotany.

A key capability in plant research is spectral phenotyping, which involves using hyperspectral imaging to capture detailed information about plant health and stress responses across a wide range of wavelengths. This technology enables the measurement of critical plant traits, such as photosynthetic activity and biochemical composition, through analysis of specific spectral bands. It can also identify stress indicators like drought or nutrient deficiencies by detecting subtle changes in spectral reflectance patterns. In EcoFABs, hyperspectral imaging is particularly powerful because it leverages the controlled environment to study the effects of induced stressors, linking lab-based observations to broader ecological or field-scale imaging data.

Segmentation is a critical step in analyzing hyperspectral plant data, allowing researchers to isolate regions of interest and extract meaningful insights from high-dimensional datasets. However, many segmentation methods rely on extensive annotated training datasets, which are often impractical to generate in dynamic and variable environments. Changes in lighting conditions, experimental setups, or hardware configurations, as well as the introduction of new plants or model systems, make it challenging to create segmentation frameworks that generalize across setups. To address this, we utilize sparse random mixed-scale networks combined with an ensemble approach, enabling robust segmentation of hyperspectral data, even under sparse training data conditions. By aggregating outputs from diverse independent classifiers, our method produces robust segmentation results while minimizing the need for labor-intensive annotation. This economical approach ensures adaptability and efficiency, even under variable conditions. The hyperspectral imaging pipeline for EcoFABs generates substantial amounts of high-dimensional data, requiring high-performance computing (HPC) resources for efficient processing, analysis, and storage. A single EcoFAB image contains 512 *×* 512 spatial pixels with 204 spectral bands - from 397 to 1000 nm - and even a small-scale experiment with 40 plants, imaged from three angles every other day for 4 weeks, can yield approximately 650 GB of data. Larger-scale, longer-term experiments with more fine-grained imaging schedules easily scale to tens of terabytes. These data volumes combined with the high-dimensional nature of the data further emphasize the need for robust, computing infrastructure. HPC enables rapid segmentation, image alignment, and spectral data processing, as well as the integration of machine learning models for phenotyping and stress detection.

In this communication, we present an above-ground image analytics pipeline tailored for EcoFABs, focusing on hyperspectral image segmentation and analysis. We detail a workflow that minimizes annotation requirements through ensemble methods and sparse random mixed-scale networks. We also address data quality assurance through image alignment and demonstrate how the results can be used to monitor plant health. The integration of innovative methods and HPC computational tools establishes a framework for bridging controlled experiments in EcoFABs with real-world agricultural and ecological applications.

## 2 Material and Methods

### 2.1 Samples & Imaging Setup

*Brachypodium distachyon*, a model grass plant, serves as the primary system for this study, leveraging its small size and genetic tractability (Novak et al., 2024). This species has emerged as an ideal candidate for EcoFAB experiments, demonstrating reproducible growth patterns and well-defined responses under external perturbations. These characteristics make it particularly suitable for studying plant-microbe interactions in controlled environments like EcoFABs, enabling standardized and reproducible experimental setups.

Hyperspectral imaging is a central tool for analyzing plant phenotypes, widely used in both field studies via drones or satellites and in specialized phenotyping facilities. In this study, we employ a Specim IQ hyperspectral camera (Behmann et al., 2018) mounted on a custom-designed motorized actuator and paired with a turntable module. This setup allows imaging from multiple perspectives, including top-down, side, and front views, capturing comprehensive data on plant structure and physiology throughout the growth cycle. Illumination is provided by a combination of LED spotlights and a custom-designed light source that integrates halogen and LED elements, ensuring uniform and extensive spectral coverage essential for accurate hyperspectral data collection.

The Specim IQ camera utilized in this study operates within a spectral range of 397–1000 nm, divided into 205 bins at 7 nm intervals. However, due to the emission spectrum of the light source used in the EcoFAB system, we truncate the spectral range to a range from 430 to 800 nm. This refined spectral dataset supports detailed analysis of plant health and physiological responses, making it well-suited for phenotypic research in the controlled environment of EcoFABs. This imaging setup enables high-throughput, multi-angle data collection critical for understanding the dynamics of plant growth and response in fabricated ecosystems.

### 2.2 Image Alignment

To ensure spatial consistency between hyperspectral images, we employ a cross-correlation-based alignment method in the Fourier domain. Each image is aligned to a high-quality reference image, which is generated through an iterative process. This alignment allows consistent placement of specific ecoFAB features across different experiments, thus making it easier to incorporate prior knowledge or to perform quality control of available data. The alignment begins by creating a binary mask of the target image, derived from the sum of its RGB channels. Otsu’s thresholding method is applied to this sum to partition the image into a foreground and background class. The binary mask is then aligned to the reference image using Fourier techniques:

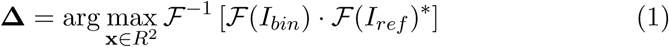

Where *I_bin_* is the binary image obtained by applying Otsu’s thresholding (Otsu, 1979) to *Σ_c_I_c_*, the sum of the RGB channels (*c*) of the target image; *I_ref_* is the reference image; *F* and *F*^−1^ denote the Fourier transform and its inverse, respectively, and **Δ** = (Δ*_x_,* Δ*_y_*) represents the offsets in the *x*- and *y*-directions needed to align the target image with the reference. This approach robustly identifies the spatial offsets that maximize the correlation between the target and reference images, ensuring accurate alignment under varying conditions.

The reference image is generated by combining 40 RGB images of plants captured in an EcoFAB across their growth cycle. These images are iteratively aligned to an initial approximate mean, and a new mean image is computed after each iteration. To enhance the quality and stability of the reference, a Gaussian blur is applied to the mean image at each step. This blurred mean serves as the alignment target for subsequent iterations. The degree of blur is gradually reduced over successive iterations, refining the alignment and producing a high-quality reference image. This final reference captures consistent structural and spatial features while minimizing noise and artifacts, providing a robust standard for aligning subsequent hyperspectral images.

Once the images are aligned, a static mask is manually generated (Sofroniew et al., 2024) to exclude regions where no plants are expected to be present. Such areas, often containing spectral signatures from reflections or other artifacts, are systematically masked out to improve the accuracy of downstream analyses. Additionally, sharp edges of the EcoFAB setup, which can distort views and introduce unwanted noise, are included in the mask to reduce false positives.

### 2.3 Network Design

For segmentation, we utilize DLSIA’s Sparse Mixed-Scale Networks (Roberts et al., 2024), a convolutional neural network architecture inspired by Mixed-Scale Dense Networks (Pelt and Sethian, 2018). SMSNets incorporate random architectures and sparse connections, leveraging stochastically configured topologies with varying random connections and convolutions of different random dilations assigned to each connection. The random nature of these model architectures produces additional diversity and higher variance among many models, making them particularly suitable for ensemble methods (Ganaie et al., 2022). By combining the efficiency of MSDNets with the benefits of randomness and sparsity, SMSNets offer a flexible and powerful tool for image segmentation tasks.

To enhance their performance for hyperspectral data, we extend these networks with a preprocessing step: rather than using the full spectral image directly in a convolution al neural network, we deploy a non-linear spectral projector, consisting of a sequence of linear projectors followed by a ReLU operator, to transform the high-dimensional hyperspectral data into a more compact representation. This latent image then serves as the input for subsequent segmentation tasks. This design offers two significant benefits. Firstly, it reduces the network’s size by minimizing the number of channels carried through inter-layer skip connections, thereby lowering computational overhead. Secondly, the compact latent representation can be easily visualized, allowing one to understand the network’s internal workings. This not only provides insights into the segmentation process but also adds a layer of transparency to the model’s decision-making. The non-linear spectral projector and the SMSNet are trained end-to-end within a single training loop, streamlining the process and ensuring optimized performance.

To enhance the robustness and reliability of segmentation, we incorporate an ensemble approach. Because the SMSNets are constructed via a stochastic network generator, every network is different and analyzes the data in a slightly different manner. By combining predictions from multiple independently trained networks, a more stable estimator is obtained, improving accuracy particularly when handling noisy or sparse datasets. Additionally, ensemble methods enable the visualization of predictor variance, offering a variability estimate for the segmentation results. This variance map is invaluable for highlighting regions of high confidence while identifying areas where predictions are uncertain or inconsistent, guiding further investigation or annotation refinements. The proposed ensemble approach provides a comprehensive solution for hyperspectral image segmentation, balancing computational efficiency, interpretability, and robustness to deliver reliable and high-quality results in the presence of minimally annotated data.

### 2.4 Training & Inference

Our training and inference approach leverages the *qlty* library to extract patches from sparsely annotated data, maximizing the utility of available training samples (Zwart, 2024b). During training, qlty is used for a sliding-window augmentation approach to enhance the diversity and quantity of our training data. This technique systematically shifts a window across the image, creating overlapping patches that expose the network to varied perspectives of the same features.

To ensure economic use of computational resources, training patches that do not contain any labeled pixels are discarded, and labels that are too close to the patch edges are removed. This approach does not only enhances the quality of the training data but also addresses the challenge of memory limitations associated with high-dimensional data. By working with smaller patches instead of the full dataset, we effectively manage out-of-core data processing, enabling training and inference without requiring the entire dataset to fit into memory. This method ensures scalability and robustness, particularly when handling large datasets in resource-constrained environments.

### 2.5 Spectral Analysis

To analyze the spectral characteristics of the plants after segmentation, we normalize the segmented plant hyperspectral data to have zero mean and unit standard deviation. This normalization ensures that the spectral data is unbiased by intensity variations - potentially due to anisotropic reflectance, shadows caused by partial occlusions and other effects - enabling a consistent and meaningful comparison across the plant.

Once images are segmented, we use Uniform Manifold Approximation and Projection (UMAP) to embed the normalized spectra into a two-dimensional space. This technique provides a visualization of the spectral data manifold, providing insights in the spectral characteristics of every plant pixel. By observing the trajectories of pixel clusters across the UMAP manifold, we can identify patterns and transitions in spectral characteristics, offering a detailed view of the plant’s physiological changes at a fine-grained spatial resolution. This approach complements an analysis using a vegetation index like the Normalized Difference Vegetation Index (NDVI), as it provides a more subtle, whole-spectrum based analysis of plant components that might otherwise be overlooked (Bannari et al., 1995).

## 3 Results & Discussion

### 3.1 Data Organization

The plant growth time series are organized in a bespoke python object, combining RGB and hyperspectral data across measured points in time. This object leverages the *zarr* storage format, which is beneficial for handling large data sets encompassing various modalities. In addition to placeholders for the actual imaging data, associated metadata can be stored as well.

### 3.2 Reference Image Construction

Using the procedure outlined above, reference images were constructed over 20 iterations by gradually reducing the blur factor, refining alignment with each step. Analysis of independent images revealed a median shift below a pixel, demonstrating the reliability of the hardware in maintaining consistent sample placement. While the segmentation networks used in this study are inherently shift-invariant, the associated static masks are not, necessitating precise alignment to ensure accurate downstream processing.

Consistent registration of the EcoFAB within the frame of reference also facilitates advanced analyses, such as the fixed localization of the stem, which is constrained by the EcoFAB hardware. This precise alignment can aid in identifying specific plant regions, enabling more detailed phenotypic studies. Additionally, the registration process serves a critical quality control role, allowing the detection of potential issues such as stalled motors and camera issues. By ensuring alignment and consistency across datasets, this registration approach enhances the robustness and reliability of the hyperspectral imaging pipeline.

### 3.3 Training

A training dataset for segmentation consisted of sparse, manually annotated data, covering 5.7% of 21 images, each of size 512 × 512 pixels. This resulted in approximately 21,000 plant pixels and 300,000 non-plant pixels. Manual annotation was performed using napari (Sofroniew et al., 2024, 2022), and due to the sparse nature of the labeling requirements, this took less then 20 minutes. To enhance the dataset, we used sliding window augmentation via the qlty library with a window size of 256 × 256 pixels and a step size of 64 × 64 pixels. Images without any labels were discarded, and labels closer than 32 pixels to the border of these patches were removed. This process increased the number of label-containing pixels in the final dataset by a factor of 43. 50% of the patches were set aside for validation purposes, driving a post-hoc early stopping criterion, where the network with the best validation score was retained at the end of 150 epochs to improve generalization (Zhang et al., 2017).

The networks contains both a spectral compressor for reducing the dimension of the data and a convolutional component that handles spatial context. The spectral compressor used a fully connected network with a channel progression of 64 → 32 → 4, including 25% dropout during training. Reducing the latent dimension of the spectral projector below 3 resulted in a decreased performance, and was fixed at 4. For the spatial component of the network, we used a Sparse Mixed Scale convolutional neural network (Roberts et al., 2024). The hyperparameters that govern the layout of these networks, the depth-vs-breath control parameter *α*, and the degree distribution parameter *γ* were chosen to balance the effective depth and width and complexity of the network and were both fixed at 0.5. Dilation factors, allowing the networks to explore scale-space in an efficient manner (Yu and Koltun, 2015), were randomly assigned to each convolutional operator between 1 and 5, inclusive. A coarse scan of the number of nodes in the graph showed that while 15 nodes yielded reasonable results, increasing it to 25 provided marginal but significant improvements. These hyperparameter searches were not exhaustive but relied on trends observed in rapid runs of a single network. Due to the stochastic nature of these networks and the associated variability of the results, exhaustive parameter tuning was deemed unnecessary and impractical. Instead, the focus was on identifying a parameter set that achieved robust-enough performance rather than optimizing for absolute maximum performance. As a result, each network has between 20,000 and 40,000 parameters, which includes the initial spectral compressor. Note that a standard UNet typically contains over 1 million parameters, necessitating more training data or more advanced regularization methods (Loshchilov and Hutter, 2017).

Training was conducted using a cross-entropy loss function with class weights of 4 for plant pixels and 1 for non-plant pixels. The networks were trained for 150 epochs with a fixed learning rate of 0.001. On an A100 GPU, training a single network required approximately 20 minutes, using a batch size of 32. Training an ensemble of five networks was completed within 40 minutes across 4 cards. A different 50% - 50% random split for training and validation of the the full dataset was used for each network, increasing the diversity of the resulting networks. Due to the embarrassingly parallel nature of the ensemble approach requiring non communication between networks and data, and the moderate amount of training data, networks can be independently trained across different GPUs.

The training process resulted in an average macro F_1_ score of 98% on the validation set, across the ensemble of five networks, demonstrating strong performance despite the imbalance in labeled pixels and the low training volume. The macro F_1_ score is calculated as the unweighted mean of F_1_ scores for each class, ensuring equal importance for plant and non-plant pixel classifications, regardless of their imbalance (Wu and Zhou, 2017).

### 3.4 Segmentation

Inference was performed on images using a patch size of 256 × 256 pixels with a step size of 248 × 248 pixels, resulting in an overlap of 8 pixels between neighboring tiles. The patches were stitched back together using the *qlty* library, yielding a total of 108 patches for a full 12 time-point growth series. For this data, the average inference time was 1.3 seconds. Reducing the patch size to 128 × 128 pixels increased the total number of patches to 300 and marginally improved inference efficiency, reducing inference time to 0.8 seconds. Given absence of need for real-time feedback for these inference tasks, further optimization was deemed unnecessary.

As outlined in the methods section, class probabilities were averaged across overlapping patches and multiple networks and subsequently renormalized. This approach also provided a means of estimating the predictive variations of the estimated class probabilities. As shown in Figure 4, the inference results highlight plant regions by overlaying the mean class probability (red) with their standard deviation (green). The visualization shown here is an illustration of typical performance across different plants and time points, supporting the reliability of the inference pipeline.

The variation in standard deviations across neural networks demonstrates the purpose of the ensembling approach: achieve a more robust performance by “aggregating knowledge across imperfectly correlated sources” (Theisen et al., 2023). This effect is further visualized in Figure 5, where we perform Singular Value Decomposition (SVD) (Jaradat et al., 2021; Eckart and Young, 1936) to evaluate the impact of ensembling across an extensive set of 30 networks and visualize how information is transformed and distributed across the network.

For a single plant series, we computed the SVD across all spectral data and visualized the top three singular images. The same process was applied to the outputs of the spectral projector, where the outputs of each network for each pixel were concatenated into a single vector for analysis. Similarly, for the spatial networks, we concatenated the final feature maps used in the linear projector that produces unnormalized log-class probabilities and subjected this to an SVD analysis. In Figure 5, each singular coefficient image - the *U* matrix remapped to the original spatial layout - is shown along with the fractional contribution of its corresponding singular value (*p_i_*) to the total power spectrum. This analysis highlights how different components - hyperspectral data, ensembled spectral projectors, and ensembled feature maps - capture and distribute the underlying information as the image is passed through the network. The distribution of the singular values varies markedly at each stage of the network. The effective rank, a measure of the dimensionality of a dataset (Roy and Vetterli, 2007), of the spectral data alone is 23, while for the concatenated spectral projector data, this rank increases to 34. For the spatial data, it increases further to 52. This growth in effective rank shows that the total information content per pixel at each stage increases due to incorportation of contextual information from neighbouring pixels. Given that each individual neural network only has 4 channels per compressed spectral image and about 30 channels for the final feature maps, the effective ranks of the data indicates the individual networks within the ensemble provide unique perspectives on the data, not equally shared with every representation provided by other networks within the ensemble. The singular maps indicate that while the convolutional neural network’s feature maps focus primarily on differentiating plant versus non-plant regions-enriching spatial and contextual cues in progressively subtler forms (edges, low-resolution representations, and positional indicators) - the latent representation derived from the spectral projector encodes more specific plant spectral information, while suppressing the importance of non-plant spectral components in comparison to the raw spectral data.

### 3.5 Validation & Spectral Analysis

The final validation of the analytics pipeline is based on the interpretation of the insights it provides. Visual inspection of the segmentation, as shown in Figure 3, confirms the validity of the high F_1_ scores achieved during training. However, the most significant insights come from the UMAP (McInnes and Healy, 2018) embedding of the segmented spectral data (Figure 6-A1). This embedding reveals a major cluster that captures systematic variations in plant spectra. The absence of large clusters of spectra contain non-plant signatures is another indication that the proposed segmentation pipeline generalizes well across the provided data.

**Figure 1:**
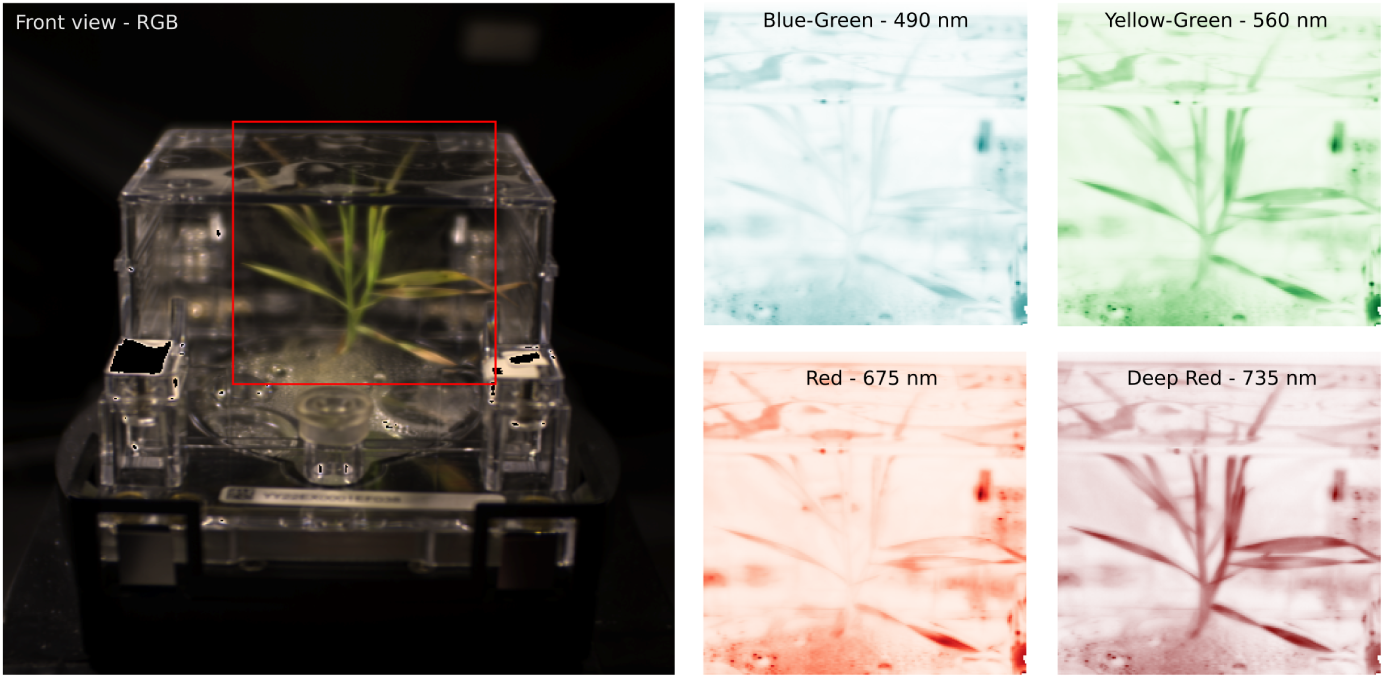
Left: A front RGB view of an ecoFAB with Brachypodium grass under stress. Right: Selected bands from the energy-resolved hyperspectral camera provide detailed reflectance observations, which can be linked to plant health and phenotypes.

**Figure 2:**
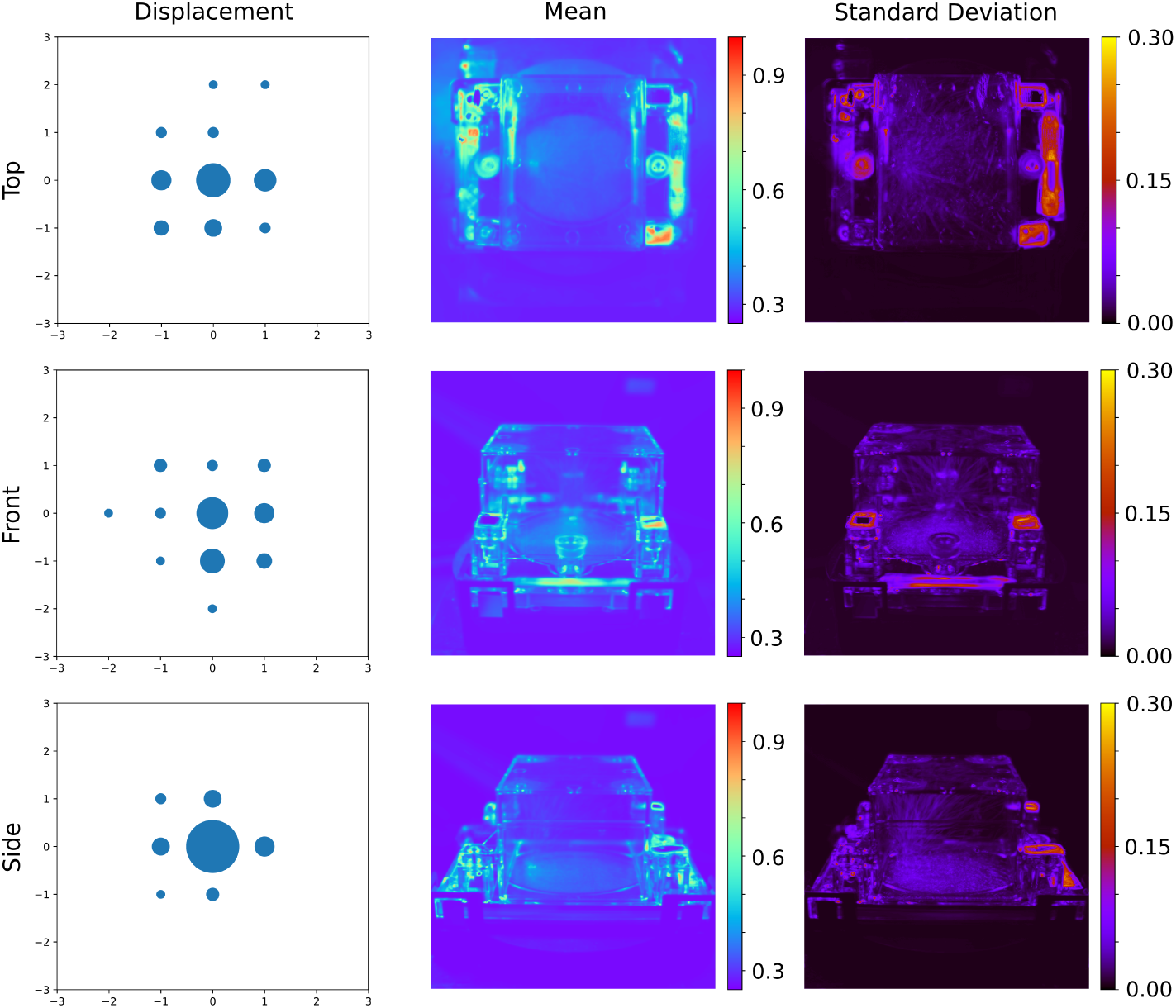
Result of the image alignment procedure. Left: Distribution of estimated shifts after aligning images to the reference image, with bubble size proportional to the number of observations with the indicated shift. Center Right: The mean and standard deviation of the shifted images aligned to a common reference, without Otsu thresholding, highlight the consistency in placement and fabrication of different ecoFABs.

**Figure 3:**
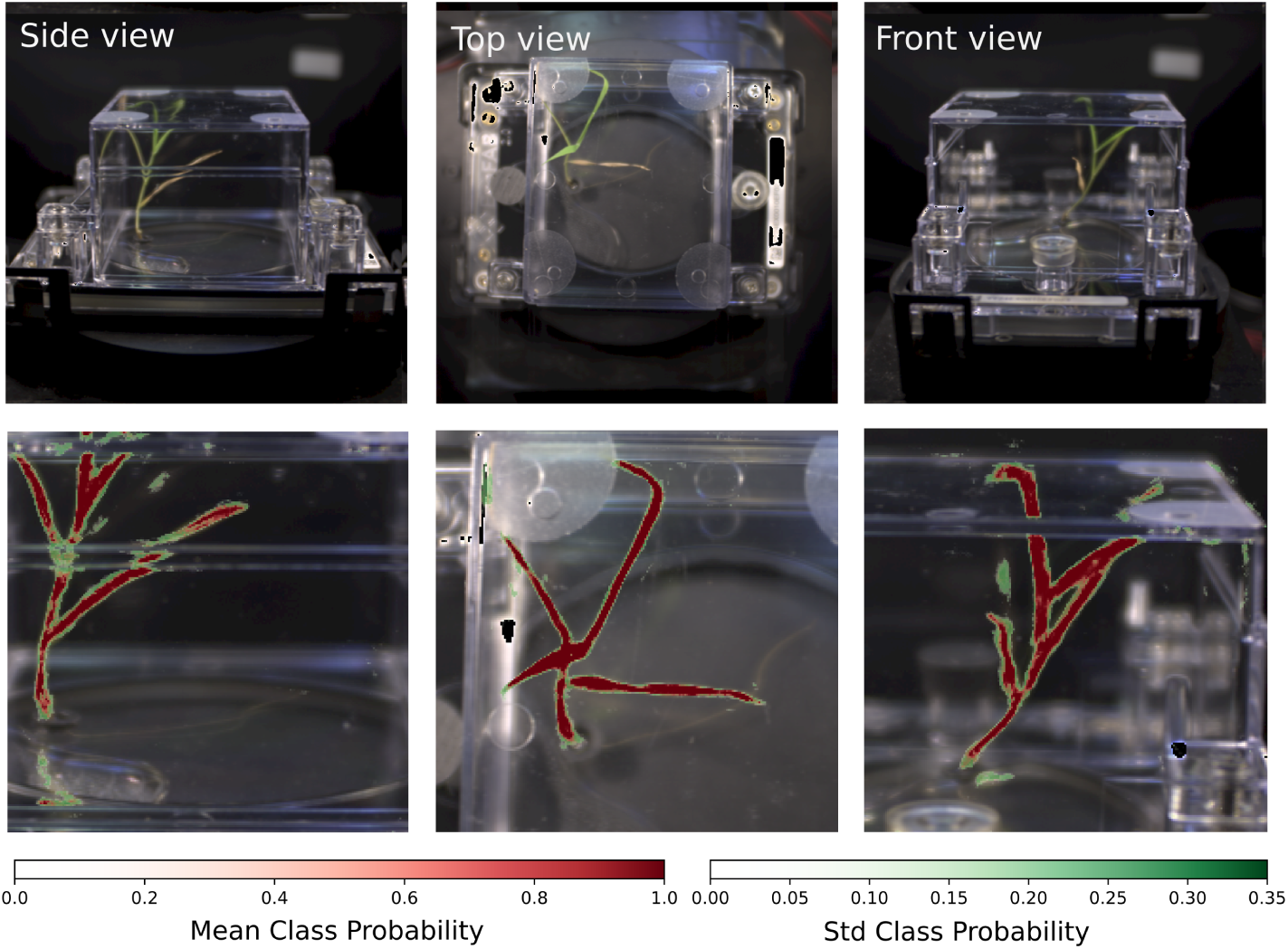
Segmentation Results. Top row: RGB images of a brachypodium grass. Bottom row: ensemble-average renormalized class probabilities and associated standard deviations based on hyperspectral data. The largest uncertainties in the segmentation are associated with the edges of the leaves as well as ecoFAB housing reflected images of the plants.

**Figure 4:**
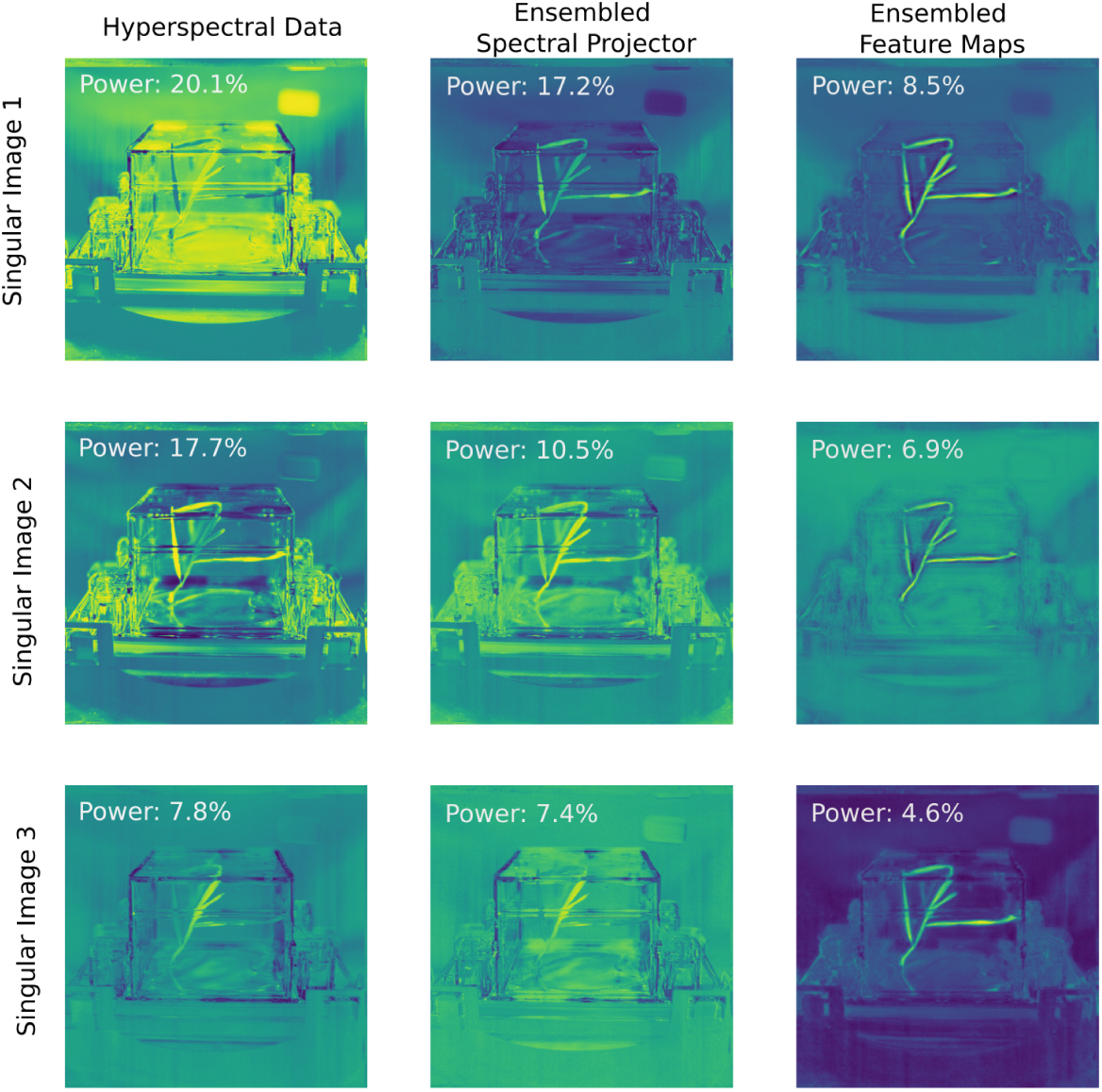
A power analysis of the hyperspectral data and the ensembled latent images indicates flatter distribution of singular values as the image progresses through the network. For the hyperspectral data without spatial context, 45.6% of the information is contained within the first 3 singular images. When contextual information is added in via the spectral projector guided by human-labeled data, the first 3 singular images contain 35.1% of the information. This number is reduced 20% when we analyze the final feature maps of the convolutional neural network. While the latter singular images mainly reveal spatial context of the pixel, the singular images of the spectral projector show a suppression of non-plant spectral features as compared to the raw data.

**Figure 5:**
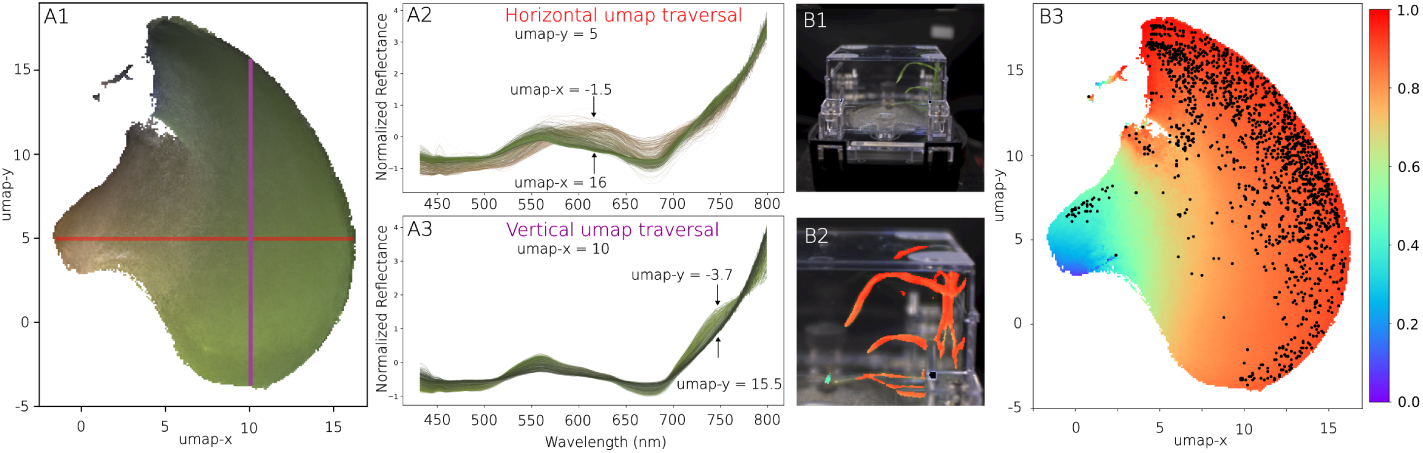
Spectral analysis. A1-3: the learned UMAP manifold of segmented plant spectra provides insights in the the type of spectra observed and their respective similarities. The color of the manifold correpsonds to the average RGB color observed in the images. A2: When traversing the manifold horizontally, the spectra transform from healthy, green leaves to stressed, brown specimen. A3: a vertical traversal across the manifold is associated with changes in chlorophyl concentration. B1: Segmentation target of a ecoFAB. B2: details of segmentation results, with plant pixels colored by their estimated NDVI. B3: Segmented plant pixels are displayed within the UMAP manifold, now colored by the NDVI, highlighting the distribution of plant health pixels throughout the plant.

The UMAP manifold representation uncovers spectral trends consistent with biological processes. For instance, traversing the horizontal and vertical axes of the UMAP embedding corresponds to changes in chlorophyll abundance and spectral characteristics indicative of senescence. The color of the manifold represents the RGB camera data, with similar colors used to highlight individual spectra. While the embedding encodes both the hyper-spectral range and RGB data, only the hyperspectral spectrum was used for distance calculations in UMAP. Importantly, the spectral analysis leverages an expanded spectral range of 430 to 800 nm, surpassing the 450 to 650 nm range used during segmentation. This expanded range is made possible by iterative improvements to the instrumentation, showcasing both the added value of hardware upgrades and the robustness of the segmentation pipeline across different hardware configurations. The enhanced spectral coverage highlights the broader utility of the analytics pipeline, enabling a more comprehensive understanding of plant spectral characteristics, as reflected in the segmentation and spectral manifold presented in the figure. The extended spectral range can for instance be used to compute the so-called Normalized Difference Vegetation Index, a single numerical indicator indicative of plant health. These trends validate the pipeline’s ability to capture meaningful spectral variations beyond simple segmentation, and can for instance be used to map out spectral components seen across a single plant, figure 6B.

## 4 Conclusions

In this study, we present a robust and scalable pipeline for hyperspectral segmentation and spectral analysis of plants in fabricated ecosystems, exemplified by EcoFABs. By leveraging Sparse Mixed-Scale Networks (SMSNets), ensemble methods, and data alignment techniques, we have demonstrated an efficient workflow capable of achieving high segmentation accuracy with minimal labeled data. The integration of advanced machine learning approaches with precise imaging setups allows us to capture meaningful spatial and spectral insights into plant physiology.

The ensemble nature of the method requires not only training a number of independent networks, but also their independent evaluation. Powerful compute resources, such as provide by HPC centers can assist in making these robust method a practical choice for a large number of scientific applications.

A key strength of the methods developed here is their broad applicability to other multidimensional imaging techniques, such as mass spectrometry imaging Velčkovć et al. (2024); Rübel et al. (2013), X-ray Fluoresence Imaging (XRF) imaging (Edwards et al., 2018), Scanning Electron Microscopy with Energy Dispersive X-ray Spectrosopy (SEM-EDX) (Rades et al., 2014) or any modality involving high-dimensional data. By combining a spectral projector coupled to a neural network, we achieve two critical outcomes: first, the SMSNet ensures robust training, effective information flow, and under reduced-data requirements, enhancing the overall efficiency of the workflow. Second, the spectral projector, trained end-to-end alongside the convolutional network, addresses the challenges of high-dimensional input data by reducing the memory footprint and learning capacity burden on the network. This enables the pipeline to process high-dimensional data more effectively, embedding context-specific information into the spectral representations.

Moreover, the spectral projector introduces non-linear effects that are guided by the segmentation outcomes, allowing the network to capture spectral patterns and features directly relevant to the task. This approach results in increased rank of the processed data matrices, embedding richer contextual information across the singular values, which enables better representation of the underlying spectral data. While dimensionality reduction techniques like SVD could be used as a preprocessing step, they are inherently unsupervised and not influenced by human-labeled segmentation data. In contrast, the spectral projector developed here integrates domain knowledge through supervised learning, ensuring that the representation aligns with the desired segmentation outcomes.

Finally, the SVD-based analysis of the ensemble offers a valuable tool for understanding and quantifying how information is distributed throughout the network. The presented segmentation workflow has the potential to serve as a foundation for a wide range of applications beyond plant phenotyping, including imaging-based material analysis, environmental monitoring, and biomedical imaging. By embedding spectral and spatial context into the analysis, the methods presented here open new avenues for exploring and interpreting complex multidimensional datasets across diverse scientific domains.

## Author Contributions

PHZ - Conceptualization, Data curation, Formal analysis, Investigation, Methodology, Software, Validation, Visualization, Writing – original draft, Writing – review & editing; PA - Data curation, Investigation, Resources, Validation, Writing – review & editing; TN - Funding acquisition, Project administration, Supervision, Writing – review & editing;

## Data Availability Statement

The trained models and sample data are deposited in Zenodo, together with hyperspectral data for selected plants (Zwart et al., 2024). Source code can be downloaded from the ecospec repository (Zwart, 2024a).

